# Beyond straight lines: migration costs considering geography enhance tracing human genetic ancestry

**DOI:** 10.64898/2026.06.16.732259

**Authors:** Jianjun Lian, Andre Python

**Affiliations:** Center for Data Science, Zhejiang University, Hangzhou, China; International Business School, Zhejiang University, Haining, China; Centre for Human Genetics, University of Oxford, Oxford, UK; School of Medicine, Zhejiang University, Hangzhou, China

## Abstract

Reconstructing the spatio-temporal history of human genetic lineages is fundamental to understanding human evolution and population distribution. While succinct tree sequences and maximum parsimony reconstruction methods applied to large-scale genomic data have improved our ability to trace the geographic history of genetic ancestry, they have essentially relied on Euclidean distances, which ineluctably ignore opportunity costs that have shaped human mobility patterns since the earliest human migrations and settlement formations. Here we propose an approach to incorporate realistic geographical migration costs through a human movement friction surface. Using simulated data mimicking the dispersal process of human migration out of Africa, we found that, compared to the Euclidean-based benchmark (M_0_), the proposed friction-based model (M_*f*_) leads to a more accurate estimation of the geographical origin (*n* = 346, accuracy M_0_ = 0.18, _*f*_ = 0.27) and genetic flux (*n* = 30, MSE M_0_ = 0.20, M_*f*_ = 0.12) through the Mandeb corridor in the Horn of Africa. We further illustrate these findings in a case study, in which our model seems to better identify plausible human migration paths from Eurasia to the Americas by accounting for geographic factors affecting migration opportunity costs, such as the Alaska Range and Rocky Mountains that represent physical barriers that constraint migration. While important migration drivers such as climate change, technological advances, social organization, and culture remain omitted here, our work highlights the importance of explicitly accounting for geographic constraints to improve our ability to reconstruct past human mobility and, ultimately, understand the evolution of human populations.

## 1 Introduction

Genetic variation observed in present-day populations reflects the cumulative effects of dispersal, demographic processes, and spatially structured interactions among individuals over time. While efforts have been dedicated to consider these aspects in population genetics, spatial and geographical information are often overlooked [Wright, 1943, Kimura and Weiss, 1964]. The recent toolkit geographic ancestor inference algorithm (GAIA) [Grundler et al., 2025] leverages the rich genealogical information contained in genomic genealogies to estimate the geographic locations of shared ancestors among sampled individuals. Genomic genealogies are commonly represented in tree sequences and encoded in the form of ancestral recombination graphs (ARG) [Griffiths and Marjoram, 1997, Wong et al., 2024, Lewanski et al., 2024]. In summary, GAIA uses ARGs together with the geographic locations of sampled individuals to estimate where their shared ancestors may have lived in the past through a process known as maximum parsimony reconstruction (MPR). This method reconstructs ancestral locations by searching for the spatial history that minimizes the total migration cost across the genealogical trees [Swofford and Maddison, 1987, Maddison, 1995]. In its simplest implementation demonstrated in Grundler et al. [2025] (hereafter referred to as the naive method), migration is assumed to occur only between neighboring grid cells, with all movements assigned equal cost, corresponding to binary adjacency matrices. However, the optimization framework implemented in GAIA—it is based on the algorithm proposed by Sankoff and Rousseau [1975] also supports weighted adjacency matrices. This flexibility allows for more complex migration costs that vary across space and makes it possible to incorporate more realistic geographical constraints, such as mountains, deserts, and other heterogeneous landscape features, into the inference process.

While a few studies have explored the effects of landscape heterogeneity on gene flow [Manel et al., 2003, Bolliger et al., 2010], these efforts have largely been confined to either forward-time simulation frameworks [Landguth and Cushman, 2010] or single-tree phylogeographic analyses [Reimering et al., 2020]. More recent advances in genome-wide genealogy inference have enabled methods such as GAIA to reconstruct spatial ancestry across entire genomes. However, they tend to rely on simple distance-based cost functions, such as Euclidean distance, which cannot capture opportunity costs from geographic features (e.g., the presence of high mountains) that affect migration paths. Bridging this gap by integrating geographic constraints and opportunities into ARG-based methods may offer a more accurate picture of spatial evolutionary histories [Petkova et al., 2016].

In many cases, incorporating data on landscape composition into population genetic studies is expected to substantially facilitate the reconstruction of evolutionary paths [McRae, 2006, Slatkin, 1993]. In GAIA, the cost matrix is a pre-specified set of parameters that defines the cost for each edge of the tree sequence given the geographic locations of its parent and child node. Since the algorithm searches for the combination of geographic locations that minimizes the total cost, it is likely that a realistic cost matrix can improve estimation accuracy of the model. However, how and to what extent does a flexible cost matrix that accounts for geographic constraints leads to improved performance remains largely unexplored. To further assess the role of geographic constraints in improving predictive performance, we compared a proposed method that accounts for opportunity costs from geographic drivers with a benchmark model using simulated data based on simple spatial models. The findings are then illustrated using a case study intended to capture potentially realistic real-world processes.

## 2 Identifying the effect of geographical information on maximum parsimony reconstruction estimates

Tree sequences provide a compact representation of genome-wide genealogical relationships by recording how local genealogies vary along the genome due to recombination [Kelleher et al., 2018, 2019]. They explicitly preserve the branching structure connecting sampled individuals to their shared ancestors, offering a natural framework for tracing genetic lineages. To accurately reconstruct geographical locations of historical nodes in a tree sequence, an MPR estimation using the naive method extracts information primarily from the structure of the trees themselves together with the topology of the spatial network [Grundler et al., 2025]. More complex models can incorporate additional geographical information encoded in the weights of network edges, which represent heterogeneous migration costs across space. We first give an intuitive description of how geographic inference can be achieved using only information from tree structure, and then investigate the relevance of additional geographical information under various spatial models. For the sake of clarity, we use simple topologies with discrete grid cells, and simulate genetic data using Selection on Linked Mutations (SLiM, v.5.2) software [Haller et al., 2026].

We first consider a simple linear stepping-stone model consisting of 20 demes, where migration is possible only between adjacent demes (i.e., from deme 1 to deme 2, then to deme 3, and so on). Each individual migrates to one of its neighboring demes with equal probability *m*, and every deme maintains a fixed carrying capacity *K*. Initially, the population is placed exclusively in deme 1. After ten thousand years of forward-time simulation, the descendants of the initial population have colonized all 20 demes. We then sample one contemporary individual from each deme and prune the phylogenetic tree to include only the samples and their ancestors, tracing back in history until we reach the most recent common ancestor (MRCA) of all the samples. The structure of the tree is shown in figure 1, along with true (panel a) and estimated (panel b) geographical positions of each node. Since recombination causes different regions of the genome to have different genealogical histories, the full tree sequence consists of many local trees that vary along the chromosome. For simplicity, we illustrate only the local tree corresponding to a specific genomic position. Although MPR estimates tend to be biased towards the geographical center of samples, it seems that the algorithm did capture well the information from tree structure in this example, since the spatial estimation of the MRCA is closer to deme 1 than to deme 20. We hypothesize that this results from an identifiable pattern in the tree structure (figure 1c). Typically, samples that are *genetically* closer to the MRCA, in the sense that their backward-time path to the MRCA consists of less coalescent events, are also *geographically* closer to the MRCA.

**Figure 1.**
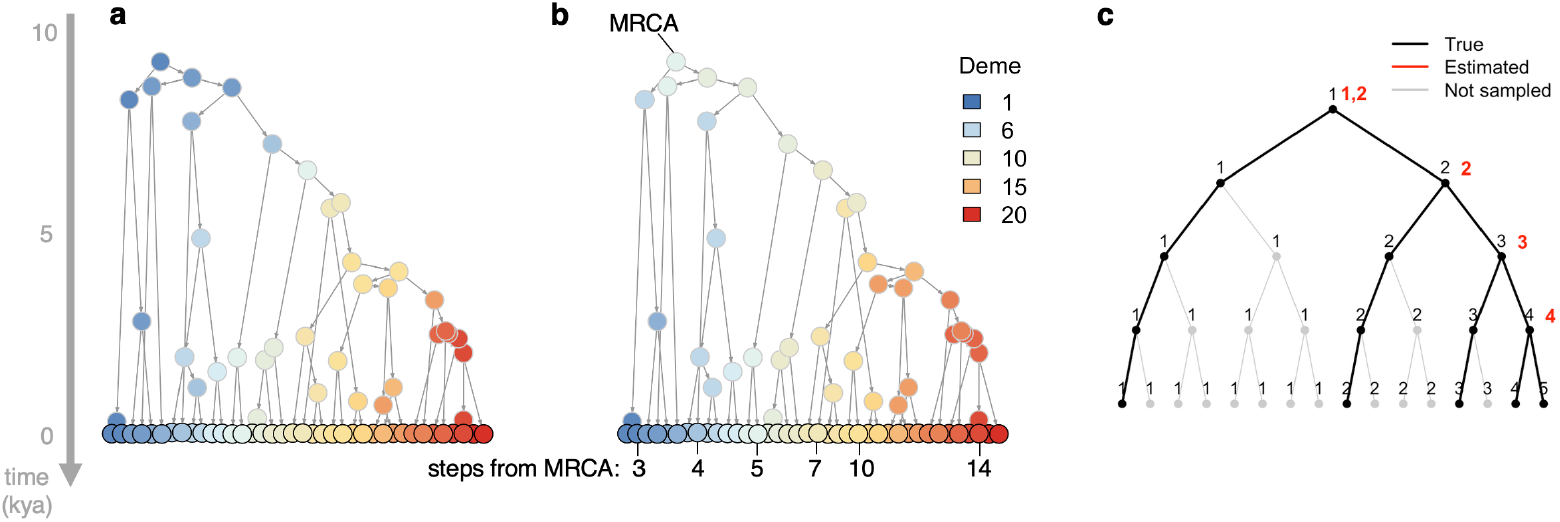
Illustration of spatial ancestry reconstruction from tree structures. (**a**) Actual geographical positions of nodes in a local tree extracted from a forward-time simulation in which an initial population from deme 1 gradually colonizes all 20 demes over 10,000 years. One contemporary individual is drawn from each deme as a sample, represented by circles with black edges. The tree is pruned to contain only the ancestry of the samples, with historical nodes represented by circles with gray edges. Geographical positions are indicated by colors, and node ages relative to the present are shown on the y-axis in thousand years (kya). (**b**) Estimated geographical positions of the same local tree inferred using the naive method applied to GAIA. Number of coalescent events needed for a sample node to trace back to the MRCA of all sample nodes on the same local tree is annotated as steps from MRCA. In general, the more steps away a sample node is from MRCA, the further it is also geographically from the position of that ancestor. (**c**) An abstracted example illustrating the characteristic relationship between tree structure and spatial history in a one-dimensional migration process from deme 1 to deme 5. True locations are shown in black and MPR estimates in red.

However, the simulation that we carried out omit important opportunity costs such as heterogeneous terrain that may affect migration in real conditions. For example, mountains and rain forests are much harder to travel across than plains. To incorporate geographic heterogeneity, we suggest to replace the binary adjacency matrix in the MPR algorithm so that migration rates may vary across the spatial network in the simulation process. If we assume full knowledge of the geographical settings, we assess whether (and if so, when) geographic information not covered by genetic information of the tree sequences may lead to a better estimation of the evolutionary tree structure. In particular, we focus on how the spatial complexity of the underlying migration space affects the usefulness of incorporating such information into the inference process.

We designed three simulations with varying spatial complexity, including a one-dimensional scenario (scenario I, see previous experiment), a two-dimensional square grid (scenario II), and a three-dimensional cubic grid (scenario III). We control the longest travel path across the network (i.e., diameter of the network) to be approximately the same for a fair comparison. We then set the following number of nodes (scenario I: 20 nodes; scenario II: side length of 10 nodes; scenario III: side length of 7 nodes). For all scenarios, we assign simple heterogeneous migration rates and runs forward-time simulations with the same population and individual settings. A few individuals are placed at a cell close to one end of the grid at initiation, and gradually fills all cells before the end of the simulation. Next we fit the simulated tree sequences with MPR algorithm using GAIA under two choices of adjacency matrix (and hence different cost matrices), leading to two distinct models: (i) *naive model (*M_0_): a cost matrix which assigns value 1 for all pairs of adjacent nodes; (ii) *friction-based model* (M_*f*_): a friction-based matrix which assigns values that are proportional to the reciprocal of the migration rate of edges to reflect that a lower migration rate between two adjacent nodes indicates higher friction values associated with a higher difficulty to travel through. We then classify all nodes of the tree sequence into four bins with equal time span and compute the average distance between estimated and true grid cell of all nodes in each bin. For a fair comparison between the two models, estimation error is evaluated using a common distance metric based on the binary adjacency structure, which counts the number of steps associated with the shortest path between the estimated cell and the true cell. Specifically, the error is defined as the minimum number of steps separating the estimated and true grid cells on the unweighted spatial network. We repeat the entire simulation-estimation pipeline for 50 rounds and perform Wilcoxon test on mean distance of the two models, as shown in figure 2.

**Figure 2.**
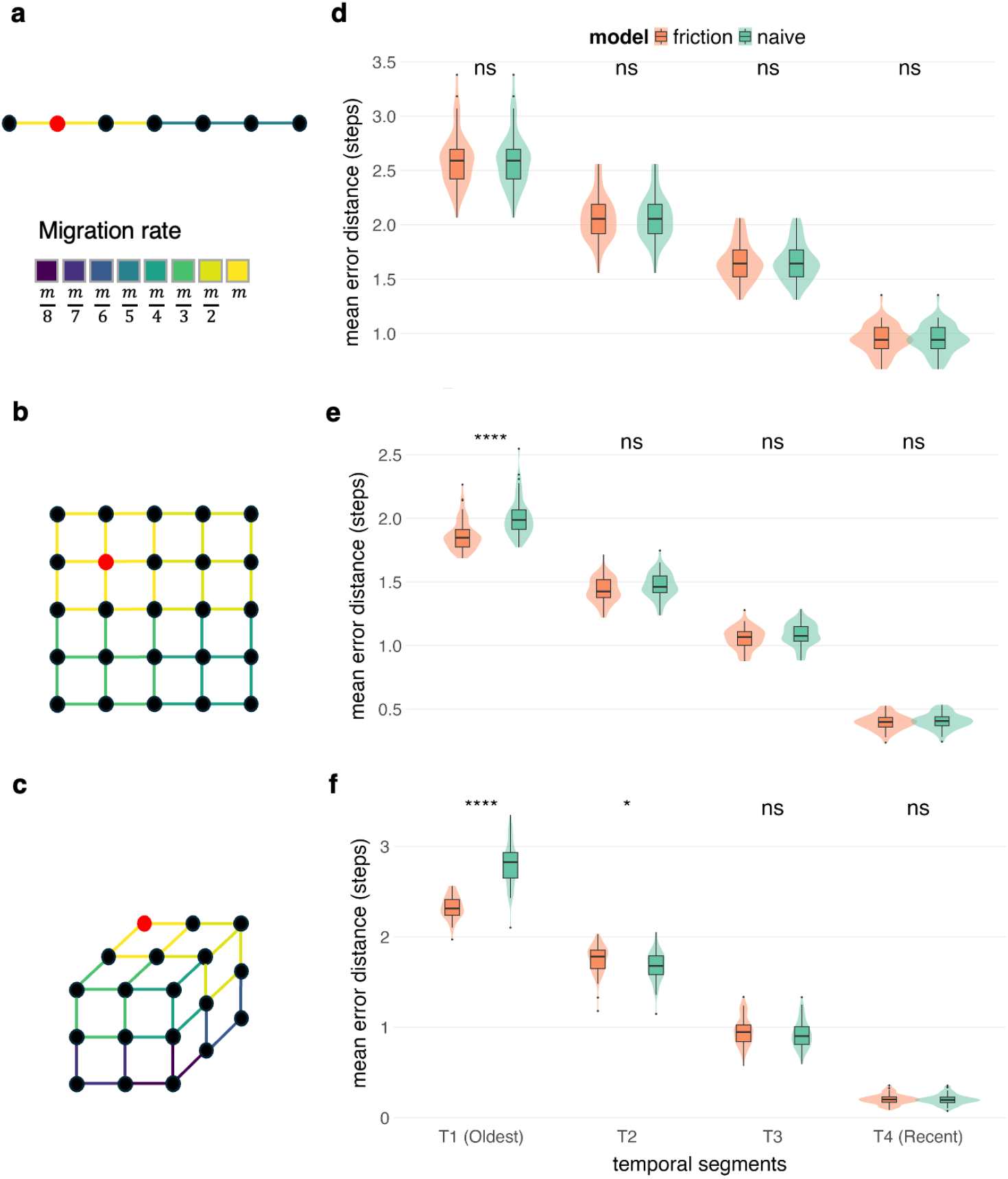
Comparison of maximum parsimony reconstruction estimation accuracy between the naive model and the friction-based model under three spatial scenarios. Simple migration models are simulated on a (**a**) one-dimensional line, a (**b**) two-dimensional square grid, and a (**c**) three-dimensional cubic grid. Note that these schematic diagrams are scaled down proportionally for clearer visualization. For each scenario, nodes represent grid cells with constant carrying capacity, and migration is allowed only through edges connecting adjacent nodes. A range of migration rates (*m*) is assigned to imitate heterogeneous landscapes, typically creating a split in half of the network across all dimensions. Edge colors represent different migration rates. The grid cell containing the initial population is shown in red, while all other cells are shown in black. Tree sequences generated from simulation of each scenario are used for MPR estimation for each model separately (naive and friction-based). The simulation-estimation pipeline is repeated for 50 rounds, and estimation accuracy is grouped according to node age in each round. Comparisons between the two models are shown for the (**d**) line, (**e**) square, and (**f**) cube scenarios with statistical significance results (ns: not significant, * *p* < 0.05, **** *p* < 10^−4^) are generated from Wilcoxon tests. The x-axis labels T1–T4 denote temporal bins ordered from the oldest ancestral nodes (T1) to the most recent nodes (T4).

The results show a common pattern of lower estimation accuracy as the nodes get deeper in history. The absolute scale of estimation error is consistent among the three scenarios. Interestingly, the estimation of the most ancient nodes via the friction-based model is statistically significantly more accurate than that of the naive model in both the square (*n* = 50, *p* = 7.05 × 10^−7^) and cube (*n* = 50, *p* = 2.42 × 10^−15^) scenarios. We also observe a larger gap in predictive performance between the native and the friction-based model at higher dimensions. We did not find evidence of differences in the line scenario (*n* = 50, *p* = 1). These results therefore indicate that an integration of migration costs may provide relevant information for the geographical estimation of ancient nodes in lattice domains of dimension two and above. An intuitive explanation is that there are more plausible paths for a piece of genetic material to travel through from the ancestor location to a specific grid cell in higher dimensions. When the choice of paths are extremely limited, the true history can easily be deduced from the tree sequence alone. This is because the structure of the tree sequence implies the merging order of contemporary nodes which, linked together, forms a path. With a large choice of paths, information from the merging order may not suffice to get the full picture; additional geographical information may matter here.

Here we quantify the concept of plausible paths using communicability, a network metric proposed by Estrada and Hatano [2008] to measure the overall strength of connection between a given pair of nodes (*i, j*) by aggregating all possible walks from *i* to *j* in a network. Given the adjacency matrix *A*, the communicability *G* between nodes *i* and *j* is defined as:

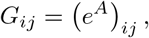

where *e*^*A*^ denotes the matrix exponential:

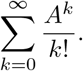

The element of the matrix (*A*^*k*^)_*ij*_ corresponds to the number of walks of length *k* from *i* to *j*. Thus, communicability can be interpreted as a weighted sum over walks of all lengths connecting the two nodes, applying weight 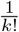 for walks with length *k*. Consequently, nodes connected by many short alternative routes have higher communicability values. This concept appears suitable for the analysis of human migration via maximum parsimony since it assumes that shorter walks are favored. To quantify the number of plausible paths in a network with *n* nodes, we propose the following metric:

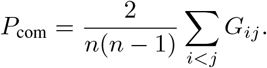

Here *P*_com_ averages *G*_*ij*_ over all unordered node pairs. To assess the differences between the friction-based and naive models, we compute the relative reduction of mean estimated error 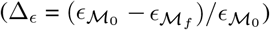 of the oldest time bin. We compute *P*_com_ for each topology reported in the main specifications (line, square, and cube).

This metric may also quantify plausible paths in more complex networks (annulus topologies in Appendix, table S1) to mimic more realistic conditions. In a context with only two paths to travel out of the ancestor location with one easier to go through than the other, the migration history through the paths can be deciphered solely from the tree sequence without information about travel constraints, even in a two-dimensional space (figure S1a). A well-documented historical case is the westward expansion and repeated migrations of nomadic and semi-nomadic groups from northern China into Central Asia between the second century BCE and the tenth century CE [Hansen, 2012, Boulnois et al., 2004, Whitfield, 2015]. In this period, human dispersal was strongly constrained to the Northern and Southern Silk Road corridors that skirted the Taklamakan Desert. Across simple and more complex topologies, our results show a systematic positive association between *P*_com_ and Δ_*ϵ*_, suggesting a better suitability of friction-based models in networks with a higher average communicability.

## 3 Tracking human ancestors with a (more) realistic migration cost

We apply the friction method to two case studies that track the history of human migration. We derive realistic migration costs from a global friction surface dataset developed by the Malaria Atlas Project. This dataset is based on the methodology introduced by Weiss et al. [2018] and further refined in Weiss et al. [2020], which provides high-resolution estimates of travel time to the nearest urban center across the global land surface. Although the dataset was originally designed to model present-day human mobility, we argue that it remains informative for reconstructing prehistoric migration patterns for two reasons. First, we use the walking-only version of the dataset, which minimizes the influence of modern transportation infrastructure such as roads and railways and therefore better reflects movement constrained primarily by natural terrain. Second, the temporal scale considered in this study is within the last 100,000 years, over which the large-scale topographic structure of the Earthsuch as major mountain ranges, continental configurations, and broad landscape barriershas remained relatively stable. While paleoclimate variation and sea-level changes certainly affected local migration opportunities, the friction surface still provides a reasonable approximation of the large-scale geographical constraints that shaped long-distance human dispersal.

We map the area of interest to a hexagonal grid, and apply mean upscaling to the friction surface to obtain the average friction within each grid cell. We define the cost of moving between a pair of adjacent cells as the time one costs to complete the travel, which is computed by multiplying the average friction between the two adjacent cells (in minutes per meter) with the distance between their center (in meter). While using real-world human genetic data may be desirable, it cannot be used to rigorously assess the validity of our model since we cannot obtain a complete and comprehensive view of the true migration paths. Thus we assess the validity of our method with data simulated to closely reflect complex migration processes that may occur in reality, and then apply it to real-world scenarios to illustrate the results.

### 3.1 Assessing the validity of our friction-based model in tracing human migration out of Africa with simulated data

We aim to verify if incorporating realistic migration costs can improve the accuracy of geographic inference in scenarios close to the real world. Using the same spatial settings as in Grundler et al. [2025], we simulate the history of human migration out of Africa across Africa and Eurasia discretized into 177 hexagonal grid cells [Barnes and Sahr, 2023], with three corridors connecting continents, including: the Strait of Gibraltar, the Sinai Peninsula, and the Strait of Mandeb [Beyin, 2011, Oppenheimer, 2012]. We use SLiM for the forward-time generation, but with different settings compared to our previous experiment. We do not apply heterogeneous migration rates that are related to travel costs in order to prevent information leakage in the estimation process. Instead, we set heterogeneous carrying capacity for each grid cell, based on fine-scale world population data obtained from Klein Goldewijk et al. [2011, 2017]. This dataset shares no common origin with the friction surface data, thus permitting no unfair advantage to the friction-based model for estimation. We aggregate global population estimates for each grid cell around 5000 BCE and apply square root scaling to derive carrying capacity used in the simulation, as shown in figure 3b. Ancestry flux is a statistical summary of the georeferenced tree sequence proposed by Grundler et al. [2025] that computes the proportion of genome of contemporary samples inherited from ancestors who moved between two adjacent cells of interest. We generated a total of 200 independent forward simulations under the same spatial setting. From these, we selected 10 representative replicates with the highest ancestry flux for each corridor (Mandeb, Sinai, and Gibraltar) for subsequent geographic ancestry inference.

**Figure 3.**
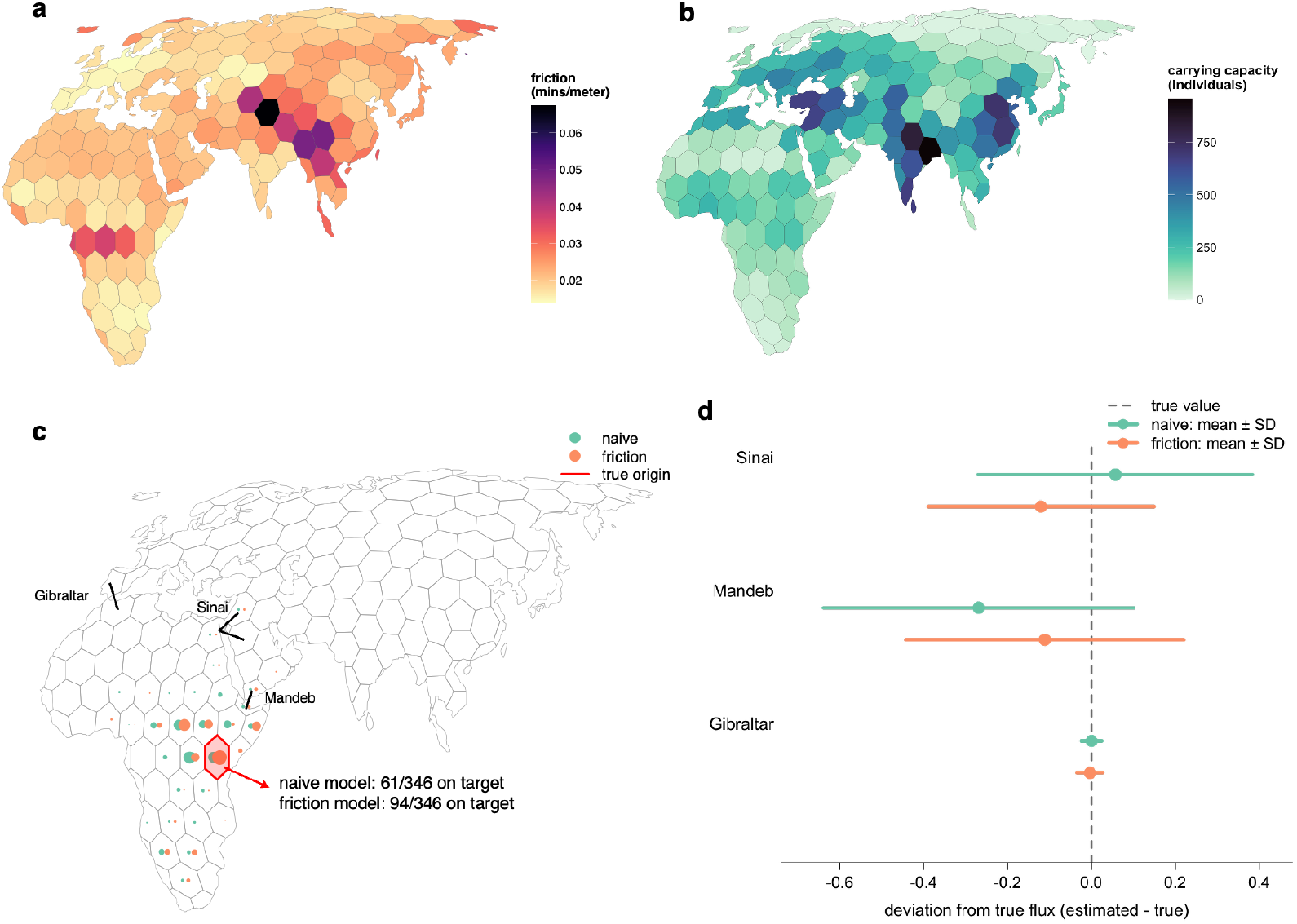
Assessing the validity of a friction-based model on simulated human migration out of Africa. **(a)** Friction surface data aggregated into hexagonal grid cells worldwide. Cells with larger friction values, i.e., harder to travel through, are represented in a darker color. **(b)** Cell-level carrying capacity mapped from fine-scale world population data estimated around 5000 B.C. Cells able to hold a larger population are illustrated with a darker color. **(c)** Estimated origin inferred from the locations of the oldest ancestors in the tree sequence across 30 replications (10 per corridor). Estimates from the naive (Euclidean-distance) model are shown in green, and those from the friction-based model in orange. Circle size reflects the number of runs in which a given cell is identified as part of the origin. The cell with red contour corresponds to the true origin. **(d)** Summary of the differences (mean ± SD) between true and modeled (naive in green, friction-based in orange) ancestry fluxes through three corridors (Gibraltar, Sinai, and Mandeb) across 30 replications.

We perform estimation using the naive and friction-based model using GAIA and compare the results. The geographical distribution of the estimation of earliest ancestors in the tree sequence is visualized in figure 3c for both models. The estimated locations from the naive model are systematically more skewed towards the west, with a lower proportion of the estimation of the origin that matches exactly the true origin compared to the friction-based model (*n* = 346, accuracy M_0_ = 0.18, M_*f*_ = 0.27). Robustness test shows that the results hold under different position of the founding deme and different simulation time length (figure S2, table S2). We also assessed the accuracy in estimating ancestry flux through the three corridors. Figure 3d shows the deviation of estimated flux from the true value, with estimation derived from the naive (green) and friction-based model (orange). Both models estimates well the flux through Gibraltar (*n* = 30, MSE M_0_ = 0.00059, M_*f*_ = 0.00093). The friction-based model performs better in estimating the flux through Sinai *n* = 30, MSE M_0_ = 0.11, M_*f*_ = 0.084), as well as for Mandeb (*n* = 30, MSE M_0_ = 0.20, M_*f*_ = 0.12). This is partly due to a better estimation of the friction-based model in scenarios where a large proportion of migrants travel through the Mandeb corridor, while the naive model constantly generates higher estimation for flux through Sinai (as acknowledged in Grundler et al. [2025]).

### 3.2 Exploratory analysis with real genetic data

We aim to provide empirical evidence of the effects of applying realistic migration costs in a case study of reconstructing human migration from Eurasia into the Americas. We discretized the spatial domain into a spatially finer hexagon grid to capture potential differences in migration patterns that may appear at a relatively fine spatial scale. We collect genomic data using a dated tree sequence inferred for chromosome 18 by Wohns et al. [2022] from the Human Genome Diversity Project [Cann et al., 2002, Li et al., 2008]. We focus on ancestry of the subset of individuals sampled from Indigenous American, Siberian, Arctic, and East Asian populations to avoid noisy signals from the extensive genomic mix of modern Americans. Our sample consists of 214 contemporary individuals from 40 populations, and the tree sequence derived from these individuals consists of 11,869 local genealogies containing 32,198 ancestral nodes and spanning over 80,000 years of human history. To balance the influence of populations while limiting the computational cost (see table S3), we perform subsampling for 100 rounds, sampling one individual from each population at a time. Within each round, the tree sequence is trimmed according to the subsamples, and locations of ancestral nodes are estimated under the naive and friction-based model using GAIA. The deduced locations are then summed together for all rounds to form a full picture of spatial population history.

Ancestors of Indigenous American populations likely originated from Northeast Asian and Siberian groups that moved into Beringia—the Ice Age land bridge between Siberia and Alaska—during the Last Glacial Maximum, approximately 26,00019,000 years before present (BP) [Goebel et al., 2008, Moreno-Mayar et al., 2018]. Archaeological evidence suggests that humans had reached South America by at least 14,500 BP [Dillehay et al., 2008]. Although the temporal accuracy of the estimates remains limited, both the naive and friction-based models successfully located the most ancient ancestors to Northeast Asia (result of the friction method shown in figure 4b). This limitation is likely due to incomplete sampling of the global genealogy. Since the analysis does not include a full tree sequence at the global scale, the algorithm is unable to accurately reconstruct migration patterns prior to approximately 26,000 BP, when the ancestors of the sampled populations would likely have coalesced with populations further west in Eurasia and Africa.

**Figure 4.**
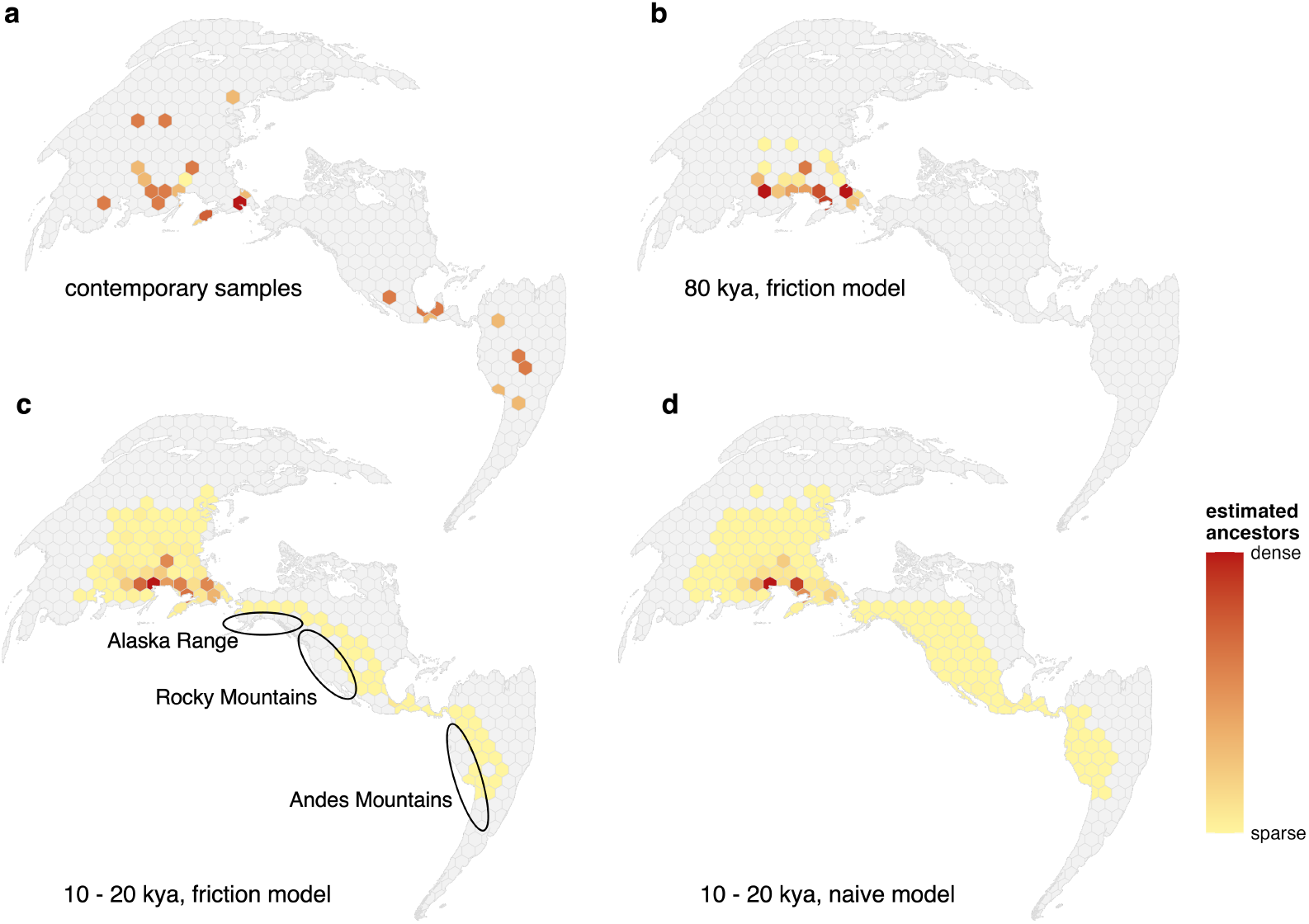
Human migration paths from Eurasia to the Americas inferred by the naive and friction-based models from 20,000 before present to now. **(a)** Spatial distribution of contemporary samples across Eurasia and the Americas. **(b)** Spatial distribution of the oldest ancestors in the tree sequences estimated from the friction-based model. **(c)** Spatial distribution of the ancestors that lived between 10,000 to 20,000 BP estimated from the friction-based model. It shows a path that connects through the three continents, avoiding major geographical barriers including the Alaska Range, the Rocky Mountains, and the Andes (black circles). **(d)** Similar plot as panel c, with the estimation changed to that of the naive model. The population distribution takes the shape of an isotropic dispersal, filling all cells that are spatially close to the shortest path in Euclidean distance. In all panels, grid cells with denser populations are colored in dark red and those with sparser populations are colored in light yellow. Grid cells without population are represented in gray.

The naive and friction-based models show consistency in the overall spatial distribution of the estimated population distribution of ancestors that lived between 10,000 to 20,000 BP (figure 4c and d). However, notice that the inferred migration from the friction-based model is consistent with migration that avoids major geographical barriers such as the Alaska Range, the Rocky Mountains, and the Andes. This is in line with common knowledge that crossing these regions is challenging and they do not offer favorable conditions for settlements. In contrast, the results from the naive model seem to generate an isotropic (i.e., directionally uniform) dispersal process that starts from the shortest path in Euclidean distance that connects the American and Eurasian samples, and systematically populates the neighboring cells along the way. The naive approach therefore appears to oversimplify the true migration process, effectively treating the landscape as an abstract two-dimensional plane with coastlines as the only geographic constraint on the habitable space.

## 4 Discussion and conclusion

We have shown that incorporating realistic geographic migration costs into maximum parsimony reconstruction can substantially improve inference of spatial genetic history, particularly in geographically complex settings. While the genealogical structure of a tree sequence already contains substantial information about historical dispersal, our results show that models accounting for realistic friction-based migration costs provide additional information that may help capture the most plausible migration routes. This effect seems to appear increasingly important as the dimensionality and connectivity of the spatial system increase. Our simulations and empirical analyses provide evidence that in some contexts, friction-based inference produces more accurate and geographically plausible ancestral reconstructions than the current benchmark that relies on Euclidean distances. These results suggest that integrating landscape heterogeneity into genealogy-based inference may provide a more realistic representation of how populations have moved through space over time.

The geographic histories inferred under our framework remain coarse approximations. Human migration is shaped by many dynamic factors that an atemporal friction surface cannot capture, such as climate change, technological innovation, social structure, and cultural behavior that change over time and across space. In addition, errors in the tree sequences introduced from upstream tools such as RELATE [Speidel et al., 2019] and TSINFER [Kelleher et al., 2019], incomplete sampling of populations, and uncertainty in ancestral recombination graphs may all propagate into geographic estimates.

Despite these limitations, incorporating realistic migration costs opens several promising directions for future research. The realism of migration cost estimation can be further improved by allowing migration costs to vary dynamically over time, capturing environmental and geographical conditions that may vary over different historical periods. Exploring how the size, shape, and resolution of the discretized domain may affect ancestral reconstruction accuracy could help establish more robust methodological guidelines for spatial genealogy inference. More broadly, combining large-scale genealogical inference with landscape-aware spatial models may provide a powerful framework for studying how geography shapes evolutionary and demographic processes through time.

## Notes

To replicate the work follow the procedure described on this GitHub link: https://github.com/SonOfNewton/geographical_ancestry_inference-paper.

## Acknowledgments

This work was supported by the National Natural Science Foundation of China (T2350610281, 82273731, and 12531013). The authors would like to thank the STAI-X 2026 organizers for creating a venue at the interface of statistics and AI, and thank the anonymous reviewers for their valuable suggestions.

## Appendix

## A Supplementary notes

### A.1 Implementation of the forward-time simulation

Forward-time simulation of all experiments are implemented in SLiM v.5.2 [Haller et al., 2026] using the same parameters as in the simulation study of Grundler et al. [2025]. Simulations were conducted under a non-Wright–Fisher framework with explicit age structure, sex, migration, and local density regulation. Individuals were considered diploid and carried a single chromosome of length 10^8^ base pairs with a recombination rate of 10^−8^ per base pair per generation. Mutations were not simulated, as only genealogical relationships were required for downstream analyses. Tree sequences were recorded throughout the simulations. Density-independent survival probability depended on age according to the following piecewise function:

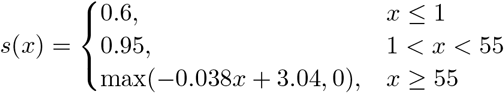

The function *s*(*x*) mimics human survival patterns and caps the age limit of every individual to 80 years. Since the number of individuals in deme *i* should be stabilized around its prespecified carrying capacity *K*_*i*_, we implement density regulation through fitness scaling. Suppose deme *i* contains *N*_*i*_ individuals with mean survival probability 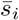, we scale the initial survival probability by a factor of 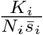. Each grid cell was modeled as a panmictic subpopulation (where all individuals are assumed to mate randomly with one another), and sexually mature individuals could migrate between neighboring subpopulations at a baseline rate of 0.001 per generation in the case studies with real-world environment. Females were reproductively active between ages 15 and 40 and could produce a single offspring per generation, while males were capable of mating between ages 15 and 55.

## A.2 Topology

For the experiments that compares the friction-based and naive methods under simple spatial networks, simulations were performed on several alternative spatial topologies. These include, in an order of ascending communicability, a line structure, three annulus lattices (from thinner to thicker grids, as described in figure S1a, b, and c), a square lattice, and a cubic lattice.

In our setting, each spatial cell corresponds to a deme connected to neighboring demes according to a predefined adjacency matrix. Base migration rate *m* takes the value 0.01 for the line scenario and 0.005 in the other scenarios. This setting guarantees similar speed of population expansion. By construction, an empty cell in a line network can only be populated with migrants from a single adjacent cell, while in higher dimensions migrants may come from multiple adjacent cells. For each scenario, we divide the network evenly based on the following migration rates: *m* and 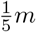 for line; *m*, 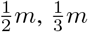 and 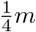 for square and annulus; and from *m* to 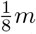 for cube. As robustness test, we also consider more complex network structures composed of annulus lattices. These annuli represent a smooth transform from a one-dimensional to a two-dimensional structure. By keeping the other parameters identical to the square scenario, we can relate differences in the predictive accuracy with the corresponding variations imposed on the network structure. The results are shown in figure S1d, e, and f, and p-values and improvements of predictive performance are summarized in table S1.

### A.3 Simulation time length

At the beginning of each simulation, a single founding deme was initialized with 10 individuals of varying ages from 0 to 60, while all remaining demes were empty. As it develops, the population gradually fills adjacent cells and occupies more area. The speed of this expansion process depends on several factors, including migration rate, population size (usually can be considered as the population capacity), and the spatial structure. In general, higher migration rate, higher population capacity of each grid cell, and higher connectivity in the sense that more cells are connected to each other (e.g., hexagon grid has a higher connectivity than square grid) are the main factors that lead to faster expansion. In the experiments using simulated genetic data, we set the parameters so as to obtain a moderate expansion speed that fits well with the simulation time length of 7,000 years. We carried out a robustness test to assess the model’s performance when part of the network may remain unpopulated at the end of the simulation, using 5,000-year period (figure S2c). We also assess the model using a 10,000-year period (figure S2d) to consider an extreme situation where an unreasonably large time length may lead to a violation of the assumptions of maximum parsimony, due to reverse migration (i.e., the local population may be shifted back and forth across the network several times) [Felsenstein, 1978, Goldberg and Igić, 2008]. The results show that accuracy of both M_0_ and M_*f*_ remains stable under all tested time lengths, with M_*f*_ constantly being 50% to 80% higher than M_0_, which indicate that the friction-based model may be more robust to variations in time length.

### A.4 Position of the founding deme

The choice of the founding deme for simulation is somewhat arbitrary and changes in its location may lead to substantial differences in the resulting migration patterns due to the heterogeneity in local environments, including landscape, population capacity, and grid structure. To assess to which extent these factors may affect the results of our comparison, we perform robustness tests with a foundation deme positioned in different locations. In the manuscript, we set the founding deme in East Africa (Tanzania) and we perform robustness tests with the founding deme located in China (figure S2a) and around Uzbekistan (figure S2b). A summary of prediction accuracy is provided in table S2. Since the three choices of founding demes have drastically different population capacity and local topological structure, the sample size varies from 346 to 793, and the degree of dispersion of the estimation also varies. However, M_*f*_ still obtains higher accuracy under all three scenarios, being 50% to 90% higher than M_0_.

## B Supplementary figures

**Figure S1.**
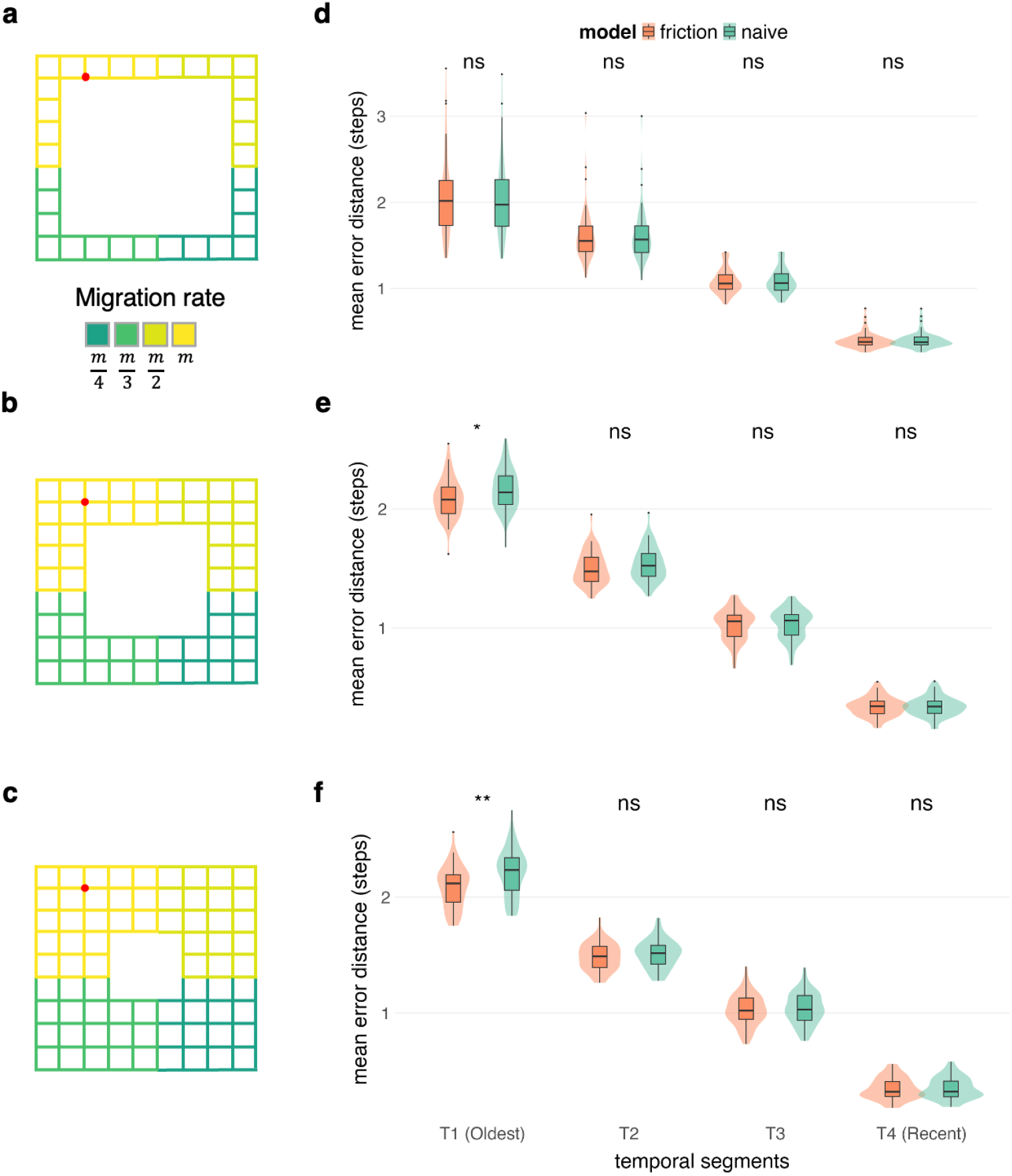
Robustness tests comparing the maximum parsimony reconstruction estimation accuracy between the naive and friction-based models in more complex network settings. We extend the network setting used in the manuscript (10 × 10 square network, figure 2b) to three additional settings with various accessibility values, by removing 6 × 6 (**a**, annulus 1), 4 × 4 (**b**, annulus 2), and 2 × 2 cells (**c**, annulus 3) leading to more complex networks. The grid cell containing the initial population is shown in red. We compared the maximum parsimony reconstruction estimation accuracy between the naive and friction-based models from *n* = 50 estimations summarized by node age. Box plots of the mean error distance (minimum number of steps separating the estimated and true grid cells) are illustrated for each model (friction-based in orange and naive in green) and scenario (**d, e, f**) and statistical significance results (ns: not significant, * *p* < 0.05, ** *p* < 0.01) are generated from Wilcoxon tests. The x-axis labels (T1T4) denote temporal bins ordered from the oldest ancestral nodes (T1) to the most recent nodes (T4).

**Figure S2.**
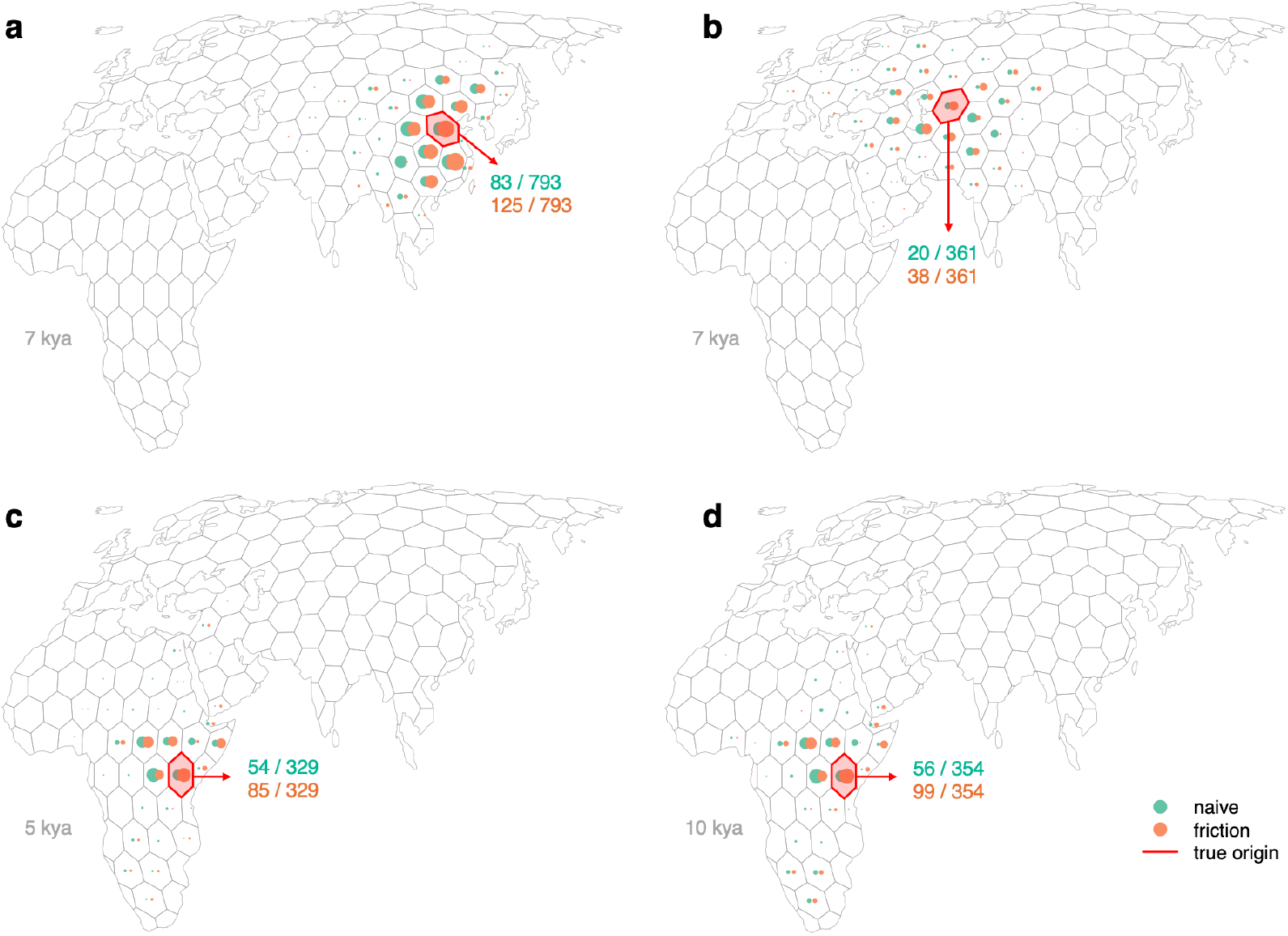
Robustness tests that vary the position of the founding deme or the simulation time length. We perform robustness test corresponding to an experiment that uses simulated genetic data and real-world environment. In the main paper, we positioned the founding deme in East Africa (around Tanzania) and the simulation runs for 7,000 years. Here we follow the same pipeline, and set the founding deme to a location in (**a**) East Asia (in China) and (**b**) Central Asia (around Uzbekistan), with simulation time length unchanged (7,000 years). In a second robustness test, the founding deme is not changed (East Africa) but we changed the simulation time length to (**c**), 5,000 years and (**d**), 10,000 years. We report for each panel the prediction accuracy of the naive and friction-based models.

## C Supplementary tables

**Table S1.** Comparison of the predictive accuracy differences between the naive and friction-based models in the oldest time bin under multiple spatial scenarios. We report the topology, number of extracted cells (if applies), dimension, communicability metric *P*_com_, mean estimated error for the naive 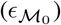 and friction-based 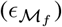 models and their relative difference, 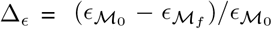 with associated p-value for each lattice topology corresponding to the results in the manuscript (line, square, and cube) and robustness tests that consider more complex network patterns (annuli with varying group of subtracted cells of 6×6, 4×4, and 2×2).

|  | main | (1) | (2) | (3) | main | main |
| --- | --- | --- | --- | --- | --- | --- |
| lattice topology | line | annulus | annulus | annulus | square | cube |
| subtracted cells | - | $6 \times 6$ | $4 \times 4$ | $2 \times 2$ | - | - |
| dimension | 1 | 2 | 2 | 2 | 2 | 3 |
| $P_{\text{com}}$ | 0.25 | 0.27 | 0.32 | 0.34 | 0.36 | 0.58 |
| $\epsilon_{\mathcal{M}_0}$ | 2.6 | 2.0 | 2.2 | 2.2 | 2.0 | 2.87 |
| $\epsilon_{\mathcal{M}_f}$ | 2.6 | 2.1 | 2.1 | 2.1 | 1.9 | 2.31 |
| $\Delta_\epsilon$ (%) | 0 | -1.0 | 3.2 | 5.4 | 6.7 | 20 |
| p-value | 1 | 0.85 | $3.4 \times 10^{-2}$ | $1.2 \times 10^{-3}$ | $7.1 \times 10^{-7}$ | $2.4 \times 10^{-15}$ |

**Table S2.** Summary of robustness tests on position of the founding deme and time length for the experiment with simulated genetic data and real-world environment. We report the accuracy (proportion of correctly predicted location of the oldest ancestry node) of the naive (M_0_) and friction-based (M_*f*_) models in alternative scenarios that vary the simulation time length including a (1) smaller time length of 5 kya and a (2) longer time length of 10 kya and the location of the founding deme in (3) China and in (4) Kazakhstan.

|  | main | (1) | (2) | (3) | (4) |
| --- | --- | --- | --- | --- | --- |
| Founding deme region | East Africa | East Africa | East Africa | East Asia | Central Asia |
| time length (kya) | 7 | 5 | 10 | 7 | 7 |
| sample size | 346 | 329 | 354 | 793 | 361 |
| accuracy $\mathcal{M}_0$ | 0.18 | 0.16 | 0.16 | 0.10 | 0.06 |
| accuracy $\mathcal{M}_f$ | **0.27** | **0.26** | **0.28** | **0.16** | **0.11** |

**Table S3.** Time complexity of MPR estimation associated with the experiment using simulated genetic data and real-world environmental conditions by sample size. The table shows the computation time per replication and in total (30 replications for each model, naive and friction-based, and assuming no parallel acceleration) using various sample sizes (number of sampled contemporary individuals) from relatively small (1) to large (6) sample sizes.

|  | (1) | (2) | (3) | (4) | (5) | (6) |
| --- | --- | --- | --- | --- | --- | --- |
| sample size | 50 | 100 | 200 | 300 | 400 | 500 |
| computation time by replication (min) | 0.20 | 0.74 | 2.7 | 5.4 | 8.5 | 12 |
| total computation time of 60 reps (min) | 12 | 44 | 162 | 324 | 510 | 720 |

## Notes

### Competing Interest Statement

The authors have declared no competing interest.

